# During natural viewing, neural processing of visual targets continues throughout saccades

**DOI:** 10.1101/2021.02.11.430486

**Authors:** Atanas D Stankov, Jonathan Touryan, Stephen Gordon, Anthony J. Ries, Jason Ki, Lucas C Parra

**Affiliations:** Biomedical Engineering Department, City College of New York, New York, NY, USA; DEVCOM Army Research Laboratory, Aberdeen Proving Ground, MD, USA; DCS Corp., Alexandria, VA, USA

**Keywords:** free viewing, natural dynamic stimuli, saccade, P300, EEG, temporal response function, TRF, active sensing

## Abstract

Relatively little is known about visual processing during free-viewing visual search in realistic dynamic environments. Free-viewing is characterized by frequent saccades. During saccades, visual processing is thought to be inhibited, yet we know that the pre-saccadic visual content can modulate post-saccadic processing. To better understand these processes in a realistic setting, we study here saccades and neural responses elicited by the appearance of visual targets in a realistic virtual environment. While subjects were being driven through a 3D virtual town they were asked to discriminate between targets that appear on the road. We found that the presence of a target enhances early occipital as well as late frontocentral saccade-related responses. The earlier potential, shortly after 125ms post-saccade onset, was enhanced for targets that appeared in peripheral vision as compared to central vision, suggesting that fast peripheral processing initiated before saccade onset. The later potential, at 195ms post-saccade onset, was strongly modulated by the visibility of the target with a spatial distribution reminiscent of the classic P300 response. Together these results suggest that, during natural viewing, neural processing of the pre-saccadic visual stimulus continues throughout the saccade, apparently unencumbered by saccadic inhibition.

## Introduction

Visual perception in the natural world requires the processing of a dynamic stimulus, with moving objects and an ever-changing perspective as the result of head and eye movements. To recreate some of this visual dynamic, recent work on visual perception uses movies and video games allowing subjects to freely scan the scene while recording brain activity and eye movements (Barczak et al., 2019; Dorr et al., 2010; Gordon et al., 2017, 2017; Konstantopoulos et al., 2010). Free-viewing is dominated by frequent saccades, which are fast eye movements that serve to update the visual input (Barczak et al., 2019; Dorr et al., 2010).

Both the saccade as well as the new visual input from the subsequent fixation produce a large post-saccadic enhancement in brain activity, which can be measured as evoked potentials on the scalp (Buonocore et al., 2020; Ehinger et al., 2015; Guérin-Dugué et al., 2018; Kazai & Yagi, 2003). Post-saccadic potentials are also known to be affected by features of the stimulus before a saccade, either in the saccade-locked event-related potentials (ERPs) or the fixation related potential (FRPs). Some of these studies use classic experimental paradigms with artificial stimuli that intentionally constrain fixations and saccades (Brouwer et al., 2013; Buonocore et al., 2020; Ehinger et al., 2015; Hiebel et al., 2018; Kazai & Yagi, 2003; Peterburs et al., 2011; Purcell et al., 2012) while others use more realistic tasks, such as reading (Dimigen et al., 2012) or natural images without restricting saccades (Devillez et al., 2015; Guérin-Dugué et al., 2018; Rämä & Baccino, 2010). Interestingly, there is also substantial evidence that, during saccades, visual processing is inhibited and subjects fail to detect changes in the visual scene or experience compressed perception (Castet & Masson, 2000; M. Ibbotson & Krekelberg, 2011; Ross et al., 1997, 2001). Conversely, after a saccade, detection performance, as well as neural activity, are enhanced as a function of pre-saccadic stimuli (Dorr & Bex, 2013; MacEvoy et al., 2008). There is also behavioral evidence for a shift of attention toward a target location starting before saccade onset (Deubel, 2008; Jonikaitis et al., 2012). Therefore, processing of the visual stimulus appears to start prior to the saccade, is inhibited during, and continues with enhancement after the saccade (M. R. Ibbotson et al., 2008). This literature spans classic experimental paradigms and free viewing of static natural images, but few of these studies deal with naturalistic tasks and natural dynamic stimuli such as video. Thus, the effect of the pre-saccadic stimulus on post-saccading processing in dynamic natural stimuli is less well understood.

We hypothesize that during a natural vision in dynamic environments there is a continuity of visual processing across a saccade. We, therefore, predicted that the visual stimulus prior to a saccade modulates post-saccading processing, as has been observed for static images. To test this we use a target discrimination task embedded in a dynamic virtual environment, while we record eye movements and EEG. Here, objects were presented on the road while subjects are being driven through a 3D virtual town (Gordon et al., 2017). They are asked to distinguish between threatening and non-threatening targets that suddenly appear in the environment. To test our predictions we analyzed saccade-evoked potentials to determine if they are affected by the presentation of targets, their visibility, and their location in the visual field. Our results suggest that neural processing of the visual stimulus, including reorienting, starts before a saccade and continues throughout the saccade without introducing delays.

## Results

Subjects (N=16) perform a target-detection task that involves discrimination of objects as a vehicle navigates through an urban environment (Fig. 1A). Participants were asked to discriminate between threatening and non-threatening targets (Fig. 1E), which appear along the road at random intervals of 1s to 3s (Fig. 1B). They respond with a button press using either left or right hand (counterbalanced across subjects). The game is presented in four different conditions: fog vs clear visibility (Fig. 1A and 1C) as well as “easy” vs “hard”. Subjects receive feedback on their discrimination performance in real-time as a numerical score on the screen, along with the scores of a virual competitor. In the “easy” condition, the virtual competitor is matched in performance, while in the “hard” condition the competitor usually outperforms the player. Each condition presents 300 targets (or “trials”) and lasts 15 minutes. During game play game we recorded 64 channels of EEG, eye movements, and button responses.

**Figure 1:**
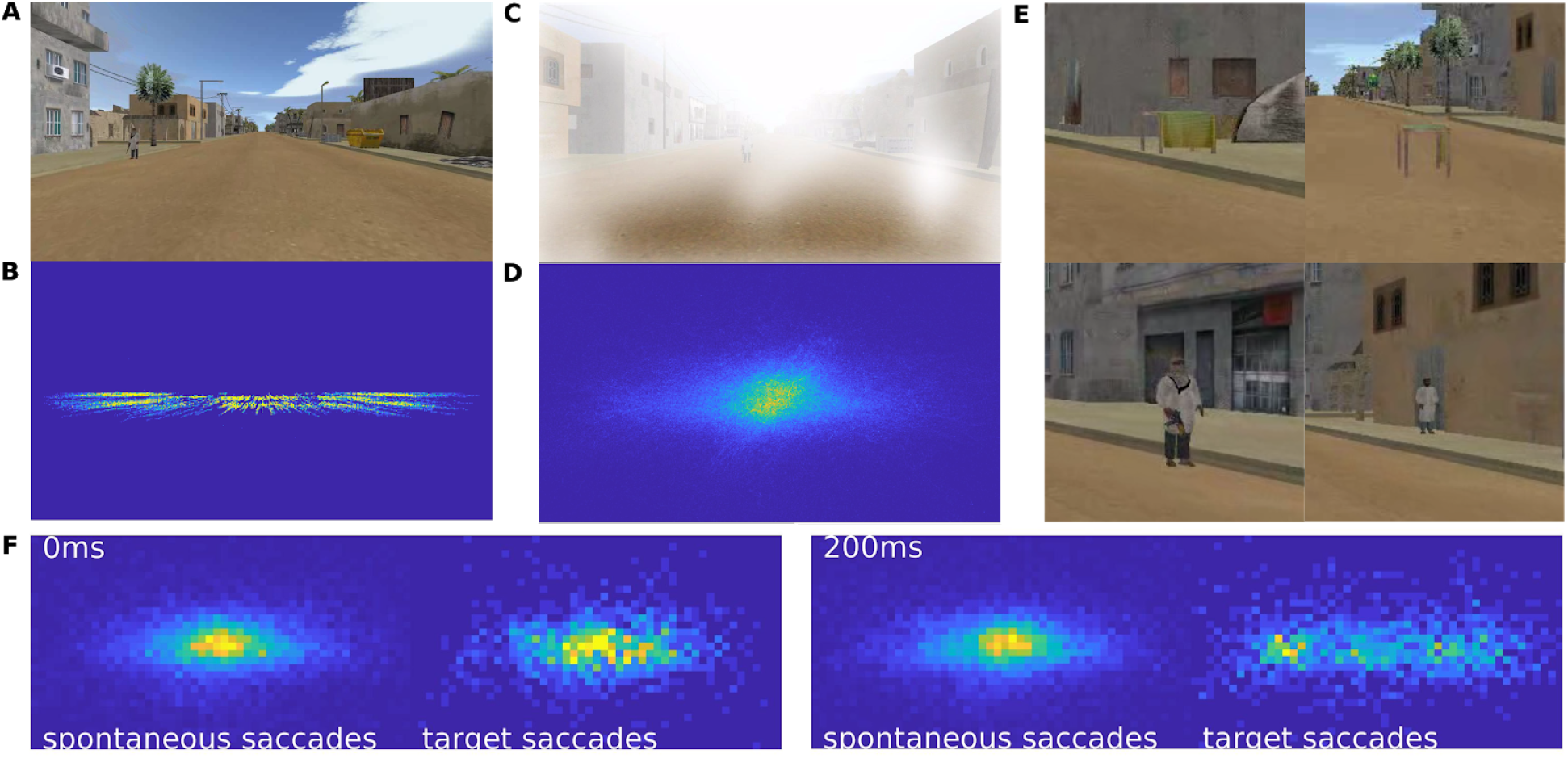
Visual environment and target discrimination task. **A:** In the discrimination task, subjects detect “targets” appear along the road in an urban environment. **B:** Distribution of target positions for 15 minutes of play. Targets move outwards as a result of forward movement of the viewer. **C:** Subjects play the game in a clear visibility condition, as well as a fog condition. **D:** Distribution of gaze position averaged over 14 subjects for 15 minutes of play. **E:** Subjects are asked to report with a button push if the target is a “threat” (left panel: covered table, armed man) or if the target is a “non-threat” (right panel: clear table or unarmed man). **F:** Eye gaze positions at 0 and 200ms post-saccade onset for spontaneous and target-evoked saccades. Target gaze goes from the center of the screen at 0ms to the left and right by 200 ms as the subjects shift their eyes to the target locations. The target-related eye gaze distribution at 200ms overlaps with the target presentation distribution as shown on panel B.

### Target presentation elicits saccades

To assess the eye movement and behavioral response during the discrimination task, we analyze gaze position, saccades, and button response times. Gaze position was focused at the center of the screen (Fig. 1D) and subjects reoriented their gaze with a saccade mostly between 100 and 450ms following target onset (Fig. 2A, blue shading). We will refer to saccades in this temporal window as “target-evoked saccades”. Target-evoked saccades tend to move gaze towards the targets, whereas spontaneous saccades do not have a preferred direction (Fig. 1F). The button responses following target presentation mainly occurred after 500 ms (Fig. 2A, dotted histograms). The timing of target-evoked saccades does not correlate with button response times (Fig. S1), suggesting that response times are not affected by the saccade.

**Figure 2:**
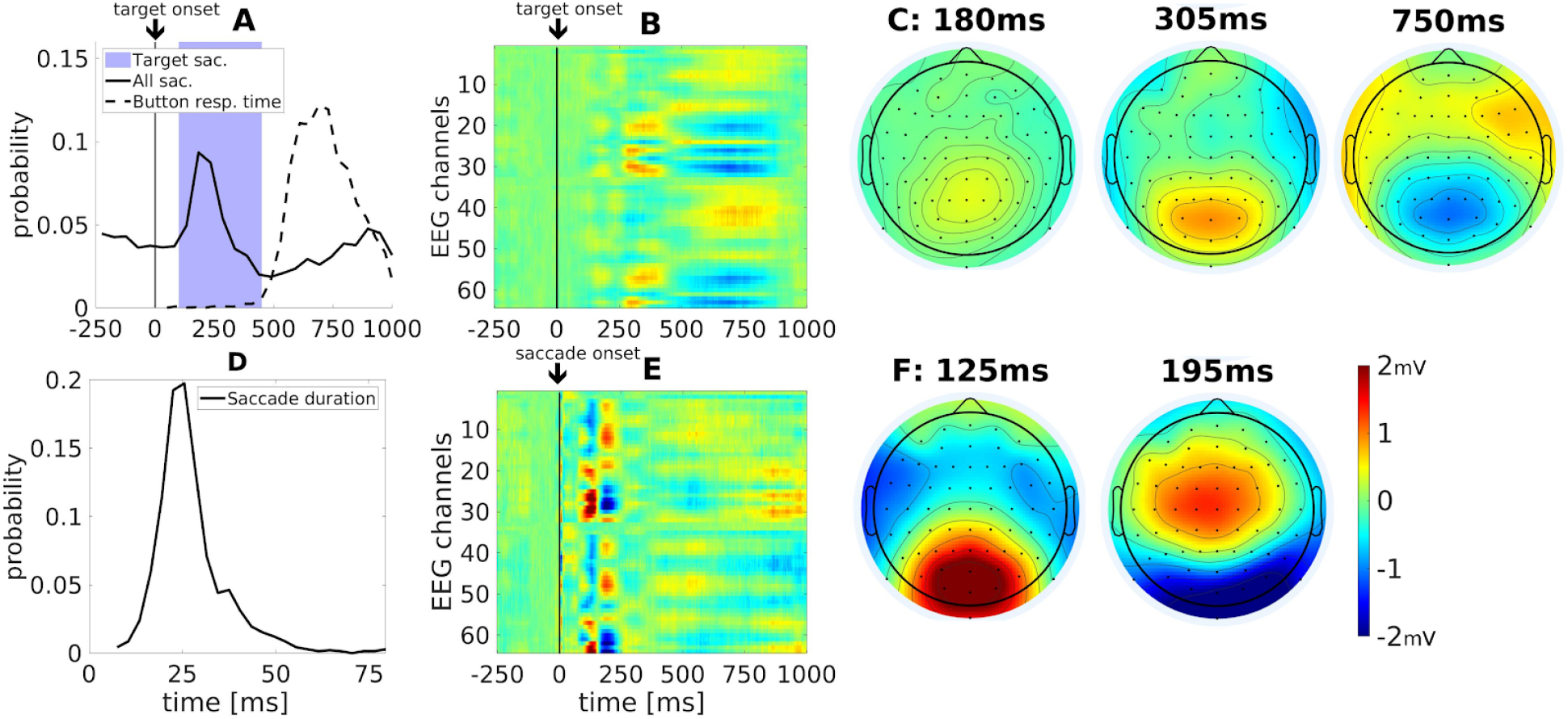
Behavioral and neural responses across all subjects and conditions for trials with a button response. **A:** Button response times (dotted line) and all detected saccades (solid line) within the same trials, which can include multiple saccades per trial. Target-evoked saccades (blue shaded time interval, 100-450ms) are selected for comparison with spontaneous saccades. **B:** Temporal response function ( TRF) locked to target presentations (N=4035). Black line at 0ms demarcates target onset. **C:** Spatial distribution of target-locked TRF at 180ms, 305ms, and 750ms. **D:** Histograms for the duration of all target evoked saccades. **E:** Saccade-related TRF for saccades following the presentation of a target (100-450ms, N=1749). Black line at 0ms demarcates saccade onset. **F:** Topographic snapshots at 125 and 195ms. Color map indicates mV identically in all panels (B, C, E and F)..

### Estimating target-evoked saccades, target-evoked potentials, and saccade-evoked potentials

Neural responses elicited during the discrimination task are characterized by multiple events, e.g. target presentation, saccades, and button responses. These events are correlated with one another (Fig. 2A) and the associated neural responses overlap in time. Therefore, simple trial averaging conventionally used to compute event related potentials can not extract the unique contributions of each of these events. Thus, we use a system identification approach to identify the individual contributions of each event to the evoked potentials (see methods). In this approach, each event contributes additively a “temporal response function” (TRF; or impulse response) to the evoked potentials. The TRFs are estimated simultaneously while taking into account correlation between different events and time points. A similar approach has been used to deconvolve overlapping responses to continuous naturalistic visual experience that involve dynamic visual input neural signals (Crosse et al., 2016; Lalor et al., 2006) and eye movements (Dandekar et al., 2011; Dimigen & Ehinger, 2021; Guérin-Dugué et al., 2018).

### Target-evoked saccades elicit unique neural response

First, we assess the effect of target-evoked saccade on evoked neural response. Here, we compute the overall TRF for each event by combining trials from all subjects and conditions. In this and all subsequent analyses we only consider trials that were followed by a button response. The TRF evoked by the target onset (Fig. 2B) reveals an initial positive potential which starts occipitally moving anteriorly to parietal electrodes in the period of 250ms-450ms (Fig. 2C). This is followed by occipital negativity and centro-frontal positivity around the time of the button push at 550ms-850ms (Fig. 2C). The initial occipital responses expected following a visual stimulus - C1 and P1 starting at 75ms (Clark et al., 1994) - are not evident here. Instead, the first target-evoked response peaks at 180ms and is more central and anterior than the conventional P1; (compare Fig. 2C to e.g. (Novitskiy et al., 2011)). Perhaps this is due to the overlapping saccades, elicited by the target onset, starting at 100ms.

The TRF for these target-evoked saccades (Fig. 2E, and as time courses in Fig. S3) resembles the temporal dynamic of the “lambda complex” (Marton et al., 1985; Yagi, 1979). However, the precise timing and spatial distribution differ from the conventional lambda complex. Here the occipital positivity peaks earlier, at 125ms, perhaps due to differences in saccade duration (Yagi, 1979). In the present study, the mean saccade duration is ~27ms with a range of durations around this mean (Fig. 2D). For these target-evoked saccades, a second TRF peak appears at 195ms, which is not evident for generic saccades, e.g. during a free-viewing image search task (Dias et al., 2013). The spatial distribution of this second peak is reminiscent of the classic P300, and specifically the more anterior P3a, which is elicited by unexpected events without the need for target discrimination (Polich, 2007). With the majority of saccades starting 200ms after target presentation, this puts the peak at 400ms, a latency similar to the P3a. Thus, both in spatial distribution, timing and task context this activity seems analogous to the P3a.

### Saccades following a target elicit stronger saccade-related TRF amplitudes

To highlight the difference in neural activity between spontaneous saccades and those evoked by the target we compared saccade-related TRFs in the presence or absence of a target. Specifically, we subtracted the TRF for target-evoked saccades (Fig. 3A) from the TRF of spontaneous saccades (Fig. 3B). There are significant differences up to 650ms following the saccade (Fig. 3C), which our system-identification approach did not attribute to the button response (see Fig. S2) that follows nearly all target-evoked saccades. Focusing on the time period before the button push (Fig. 3E), we find significant enhancement of the same two peaks at 125ms and 195ms (Fig. 3D). Early saccade-evoked potentials are known to be modulated with saccade amplitude (Yagi, 1979). Note that saccade duration did not differ between target-evoked and spontaneous saccades (Fig. 3F; t(15)=-0.38, p=0.70), which rules out saccade amplitude as a confounding factor (amplitude and duration correlate). Instead, it is the presence of the target that enhances saccade-related TRFs as early as 125 ms after saccade onset.

**Figure 3:**
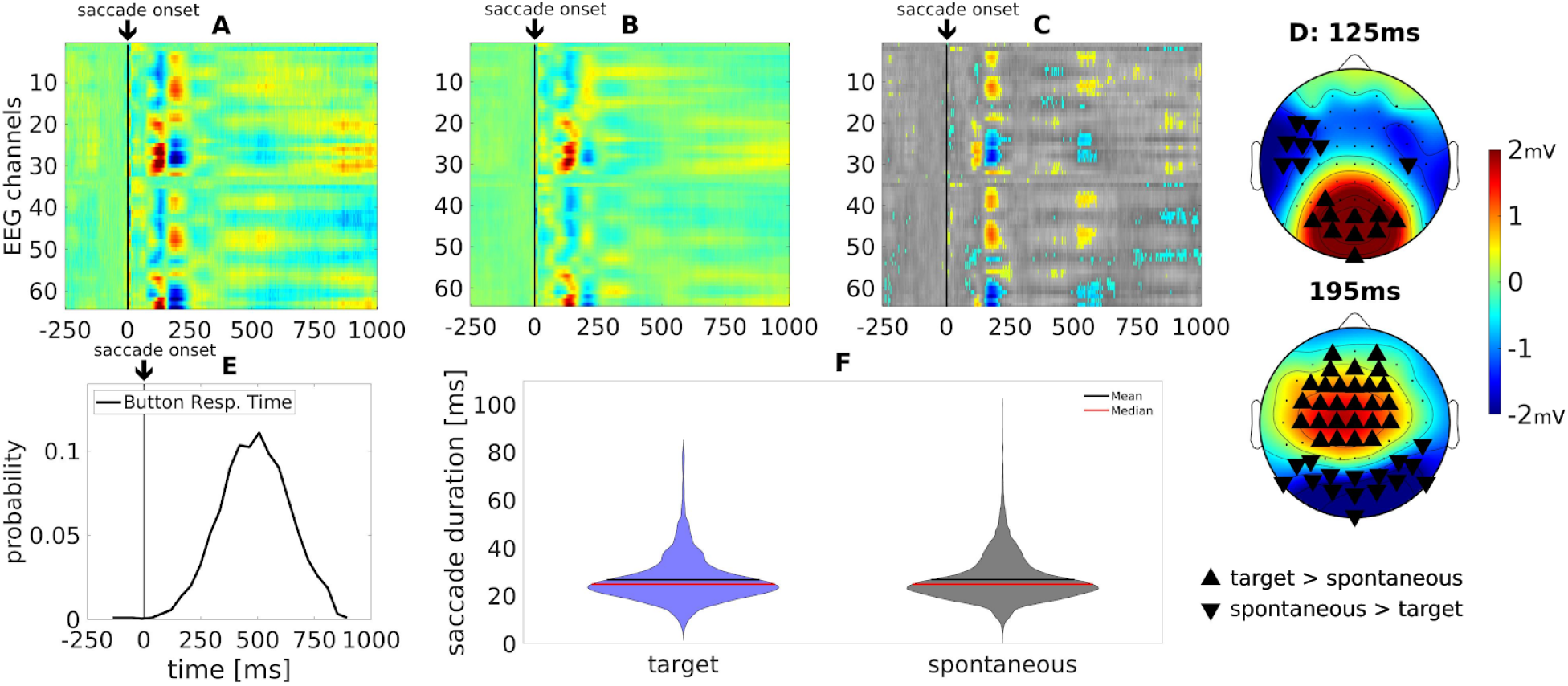
Differences in saccade-related TRFs between spontaneous and target elicited saccades. **A:** TRF for target-evoked saccades (N=1749), which are defined as saccades within 100-450ms after target presentation. **B:** TRF for spontaneous saccades (N=12437), which are those saccades not preceded by a target for at least 1000 ms. **C:** The difference in saccade-related TRF between target-evoked minus spontaneous saccades. Significant differences are shown in color, superimposed on non-significant differences in gray. Significance is computed by permuting event onsets between the two conditions 500 times. This provides a surrogate distribution at which a two-tailed significance was determined at p<0.05 after false-discovery rate correction for multiple comparisons. Color map indicates mV identically in all panels (A-D), however, in panel C color indicates voltage difference between types of saccades. **D:** Spatial distribution of the TRF difference at 125ms and 195ms. **E:** Button response times curve with respect to saccade onset. **F:** Density plots of saccade durations for target-evoked and spontaneous saccades.

### Target visibility affected late but not early saccade-related TRFs

We expected that poor visibility (low contrast targets) would result in slower button responses and weaker evoked responses as compared to the clear visibility condition (high contrast targets). In the fog condition, subjects were significantly slower in their target-evoked saccades (Fig. 4E; t(15)=3.42, p=0.0038) and button responses (Fig. 4F; t(15)=4.91, p=1.9e-4) as compared to the clear condition. To test for differences in evoked responses we subtracted the target-related TRFs of the clear visibility conditions (Fig. 4A) from that of the fog condition (Fig. 4B). We see significant differences starting at 150ms and lasting at least until 1s after target onset (Fig. 4C). As expected, differences reflect a drop in TRFs magnitude for the fog conditions, i.e. positivity increases and negativity decreases (Fig. 4D).

**Figure 4:**
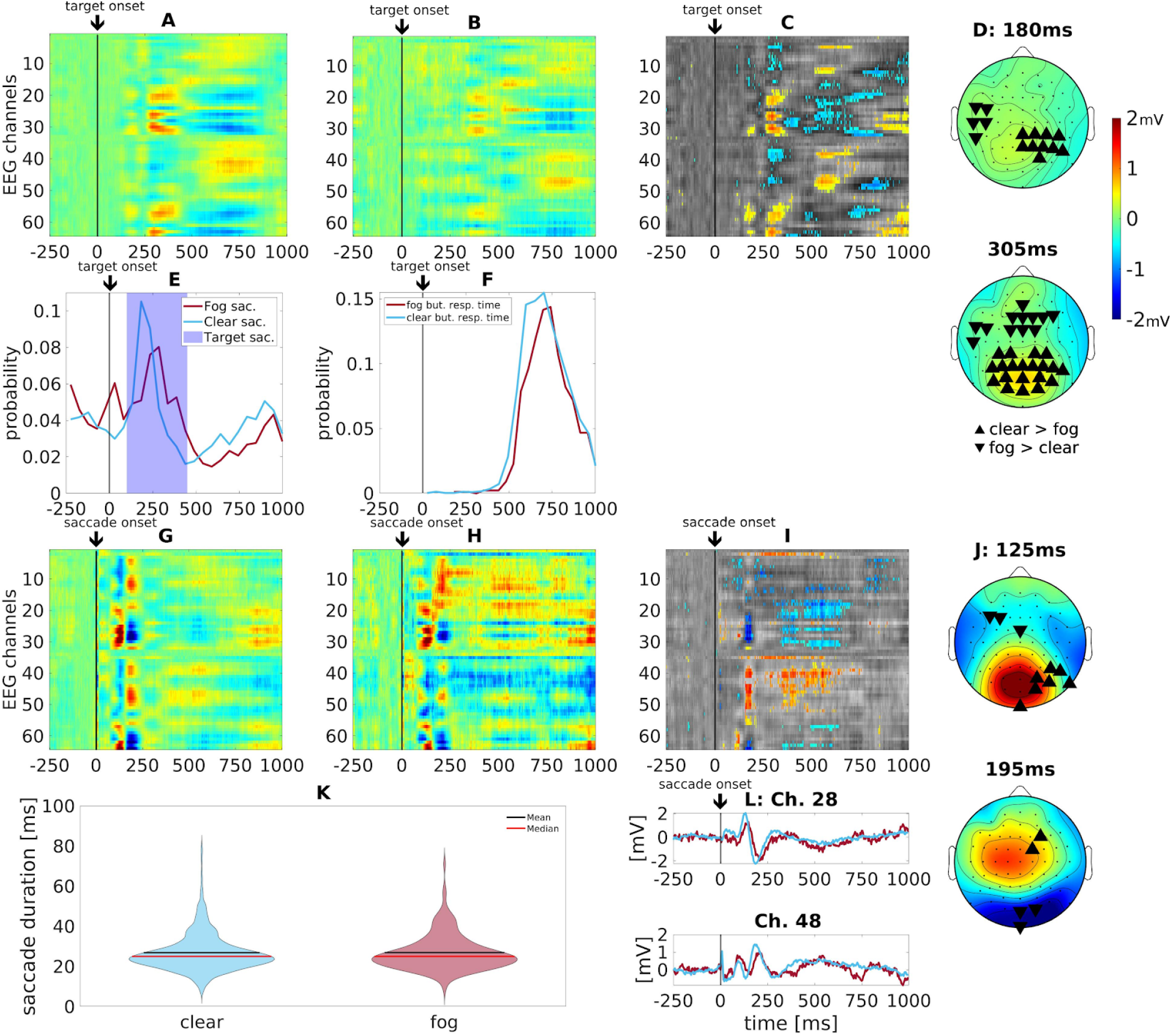
Target visibility: difference between clear and poor stimulus visibility. All trials here had button responses. **A:** Target-locked TRF in the clear visibility condition (N=3159) shows an earlier and higher amplitude response relative to the **B:** the fog condition (N=876). **C**: Differences in target-locked TRFs in the two visibility conditions (clear - fog). Significant differences are shown in color, superimposed on non-significant differences in gray. **D:** Topographic snapshots at 180ms and 305ms show the spatial distribution of the differences in panel C. **E:** Saccade histograms for fog and clear visibility conditions. Subjects saccade to target significantly later in the clear compared to the fog condition. **F:** Similarly subjects’ button press responses are also significantly later in the clear compared to the fog condition. **G:** Saccades-locked TRF to clear condition (N=1382) show a strong response but this is absent in the **H:** fog condition (N=367). **I**: Comparing the condition in the saccade-locked TRFs shows a contrast strongly driven by the saccade responses for clear visibility. Significant differences are shown in color, superimposed on non-significant differences in gray. **J:** At 125ms and 195ms this result is spatially similar to Fig. 2D. **K:** Density plots of saccade duration times for the clear and fog condition saccades. **L:** EEG traces for channels 28 and 48 showing the significance between clear (blue) and fog (red) is driven by higher amplitude and the earlier neural response of the clear condition. Color map indicates mV identically in all panels (A-D, G-J), however, in panel C and I color indicates voltage difference between visibility conditions.

Saccade-locked responses to the clear and foggy conditions both show robust response (Fig 4G and H respectively). In computing the differences for the saccade-locked potentials (Fig. 4I), we see both early and late response (125 and 195ms) are reduced in magnitude (Fig. 4J). These differences are not resulting from differences in saccade amplitude as their mean values did not differ significantly (Fig 4K; t(15)=-0.10, p= 0.92). Instead, the differences reflect a drop in scalp voltage amplitude in the fog conditions as well as a delay of the saccade-related TRF activity seen around 195ms (Fig. 4L).

### Peripheral targets elicit stronger saccade-locked TRFs, without affecting response times

We also expected that targets appearing in the periphery of vision would elicit larger saccades, and possibly differ in how the target is processed. To test for this, targets were split into equal numbers of “peripheral targets’’ and “central targets’’ depending on the distance between the subject’s gaze and the location of the target when it first appeared on the screen. The median distance for this split was 8.58° in visual angle (median established over all trials across all subjects). Subjects may have saccading earlier for central targets (Fig. 5A, 100-450ms after target onset; t(15)=-1.90, p=0.08) and had a larger number of detected saccades (Fig. 5A, t(15)=-5.39, p=8e-06). Saccade durations showed no significant difference (Fig. 5F; t(15)=-1.66, p=0.097). Additionally, there was no significant difference in mean button response times (Fig. 5E, t(15)=-1.51, p=0.14). However, they responded more frequently (Fig. 5D) and with higher accuracy to central targets (Fig. 5G, signed-rank test: z(15)=-3.10, p = 0.0019). We find significant differences in target-related TRF between the peripheral and central targets starting at 305 ms (Fig. 5B), which is shortly after the majority of saccades have terminated. We also did find differences in saccade-locked TRF at starting with saccade onset and including the activity we previously observed at 125 ms (Fig. 5E, 5F, here differences are more apparent at 129ms). In total, equal response time, as well as an effect of pre-saccade target location on evoked response suggests that processing of the target started prior to saccade onset.

**Figure 5:**
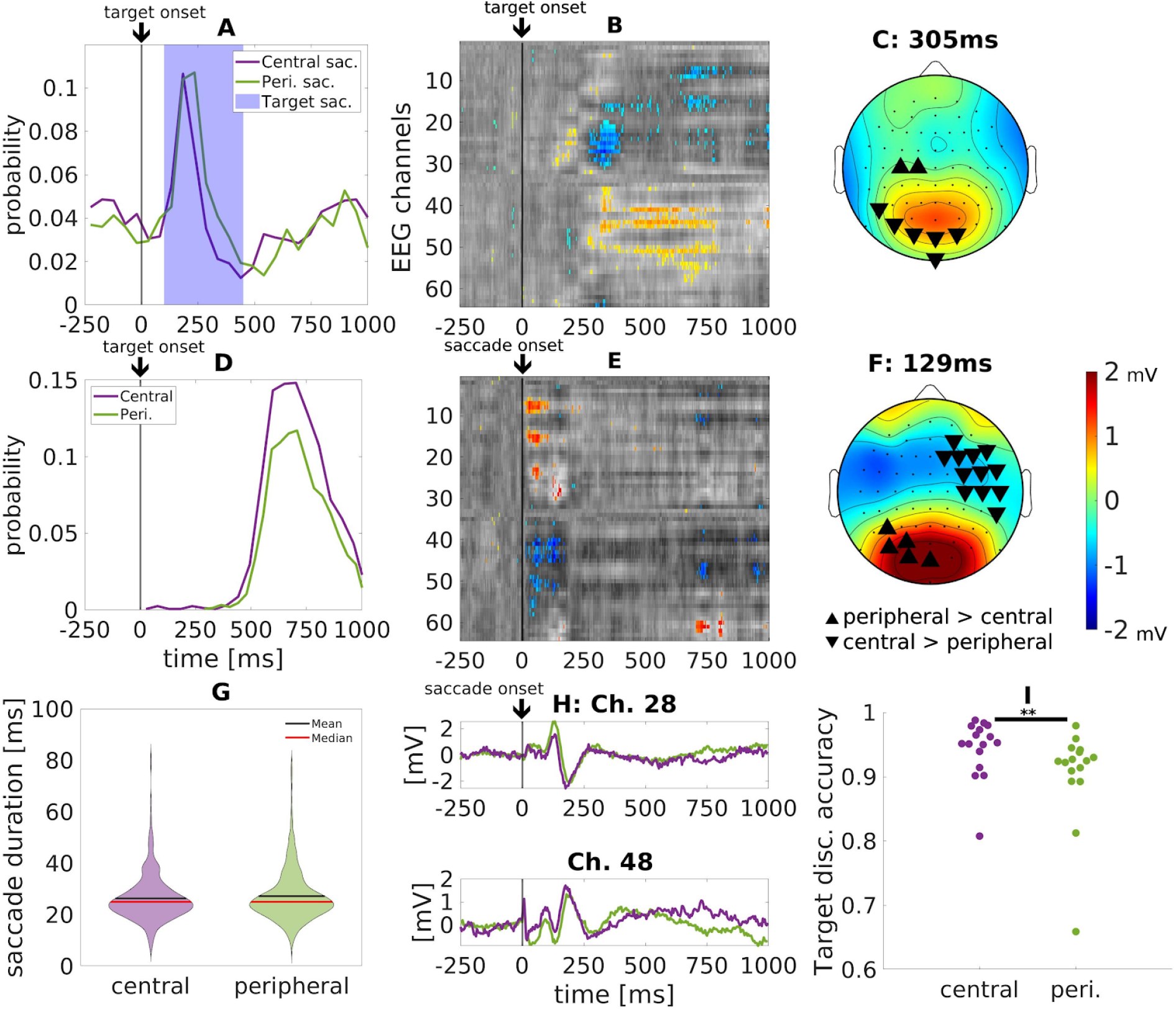
Target location: differences between responses to peripheral and central targets. **A:** Saccade time histograms relative to target onset. **B:** Difference in target-locked TRF, taking peripheral (N=1546) minus central (N=1613) target locations. Significant differences are shown in color, superimposed on non-significant differences in gray. **C:** Topographic snapshot of target-locked TRF at 305ms. **D:** Button response histogram locked to target onset for central and peripheral targets. **E:** Difference in saccade-locked TRF between peripheral (N=830) minus central (N=552) targets. **F:** Topographic snapshot of saccade-locked TRF at 129ms. **G:** Distribution of saccade duration for the central and peripheral targets (t(15)=-1.66, p=0.097). **H:** TRF for channels 28 and 48 showing that differences in E are due to differences in amplitude between peripheral (green) and central (violet) targets. **I:** Button response accuracy for central or peripheral targets. Color map indicates mV identically in all panels (B, C E, F), however, in panels B and E color indicates voltage difference between types target locations.

In closing, we want to note that the results reported in Fig. 3-5 were all computed in the “easy” game condition, and reported here only if they reproduced similarly for the “hard” game conditions (See Fig. S4-6), as no significant differences between easy and hard game conditions were found. We also did not find differences between threat and no-threat targets that were consistent across game difficulty (Fig S7).

## Discussion

To summarize, targets appearing suddenly in a dynamic visual scene elicit immediate saccades. Early activity (at 125ms) following the onset of these saccades is modulated by visibility as well as the location of the target. This suggests that visual processing of the target continues throughout the period of the ensuing saccade. Late activity (195ms) following the onset of these saccades is not present during spontaneous saccades and has a scalp distribution consistent with the classic P300. This suggests that the orienting response typically observed during static vision is not delayed by the intervening saccade.

Here we made an attempt to separate neural activity evoked by the target from neural activity associated with saccades. This is complicated by the fact that the target presentation is reliably followed by saccades. Essentially, we extract activity locked to either target onset or saccade onset, while removing correlation induced by their overlapping co-occurrence. Interestingly, the presence and properties of a target modify the activity evoked by saccades, despite having removed target-locked activity. This finding implies that the presence of a target has a continued effect on stimulus processing after saccades.

Differences in peak neural activity between target and spontaneous saccades are perhaps most evident 195ms after saccade onset. This peak has a distribution similar to the well known P300, specifically the P3a (Polich, 2007). Similar to the conventional P3a, it peaks at approximately 400ms after target onset. Remarkably, this activity does not appear with that same spatial distribution in the target evoked activity, i.e. locked to the target presentation. Yet the P300 has largely been studied in the context of target presentation and time-locked to stimulus onset. Whereas here, given its alignment to the saccade, the activity is variable in time relative to target onset. P300 has been interpreted as a marker of surprise and updating of working memory, while others have discussed it in the context of reorienting attention (Donchin, 1981; Polich, 2007). This is consistent with the present context, where subjects redirect their overt attention to the sudden appearance of the target.

There is previous evidence that the P300 occurs after a saccade onto a target (Brouwer et al., 2013; Devillez et al., 2015) and does not fundamentally differ from that found under conventional fixation conditions (Dandekar et al., 2012). Here we extend these findings to dynamic stimuli during free-viewing, where we find that the P300 is temporally coupled to the saccade, i.e. the reorienting event, rather than the target presentation itself, as in traditional constrained tasks. Yet, its timing suggests that its computation already started with the target popping up on the screen. Therefore, we favor the interpretation of P300 found here as a marker of surprise, which may play a role in modern theories of synaptic learning (Gerstner et al., 2018). In these theories, the act of selecting a stimulus plays an important role in tagging synapses (Roelfsema & Holtmaat, 2018). Therefore, during free viewing in a dynamic visual environment, the saccade-to-target may be the dominant event, rather than the appearance of a target. Regardless, our results argue for a new conception of the real-world P300 that is more directly linked with the saccades-to-target rather than the stimulus presentation under fixation, which rarely happens in real life.

We have argued for the primacy of saccade-related processing, as opposed to target-related processing. Yet, we find a clear early effect of the target stimulus on the saccade related potentials, specifically, visibility and screen location (i.e. clear and fog, central and peripheral targets). Visibility in the clear and fog conditions in our study is most closely related to contrast variations in traditional studies. The activity found at 125ms is consistent in timing and spatial distribution with early visual responses. Specifically, the C1 occipital negativity peaks 75ms after stimulus presentation (Clark et al., 1994) and its magnitude increases with stimulus contrast (Foxe et al., 2008; Gebodh et al., 2017). Note that this component is measured relative to the onset of a peripheral stimulus during fixation. Given the approximate 30ms average saccade duration in our study, the contrast dependence we see at 125ms after saccade onset can readily be explained as modulation of visual processing starting with fixation onset.

Interestingly, the spatial distribution of this post-saccadic activity is consistent with the traditional C1 response, which has occipital positivity for visual stimuli appearing in the lower visual field (Gebodh et al., 2017). This is consistent with targets appearing laterally below the mean fixation point. In contrast to the TRF associated with target-evoked saccades, the TRF associated with target appearance does not match conventional visual evoked responses, neither in timing nor spatial distribution. Thus saccade TRFs are a better match to early visual processing under fixation conditions than conventional stimulus-evoked TRFs.

While contrast dependence of the early activity may result from post-saccadic processing, the dependence on the location is harder to explain. After both peripheral and central saccades subjects are foveating the target, and therefore visual input after fixation onset is approximately the same. Therefore, the location dependence suggests that visual processing of the target starts prior to the saccade and has a delayed influence on visual processing at 129ms after the saccade. The fact that button response times are independent of where the target appears on the screen also suggests that target processing is undisturbed by the intervening saccades.

A similar picture emerges if we interpret this activity as part of the lambda complex. The lambda complex is an evoked activity found in the fixation-locked analysis. It has a dominant positive deflection over occidental electrodes after fixation, appearing at 125-140ms when the saccade locked (Yagi, 1979). For relatively short saccades of 30ms, it is expected to be at 100-125ms (Thickbroom et al., 1991). Its neural generator is thought to be the same as early occipital responses in the conventional stimulus-evoked analysis (Kazai & Yagi, 2003). Since the lambda response is known to be sensitive to the afferent sensory input at fixation (Ries et al., 2018; Yagi, 1981) it could be argued that the larger saccade-locked activity found here is due to the differences in sensory information at fixation. e.g. luminance contrast of the target may be higher than spontaneous fixation locations. However, there are saccade-locked components at earlier latencies relative to the dominant lambda complex over occipital electrodes (Thickbroom et al., 1991), and there is support for a fixation-locked component at approx 50ms that differentiates salience/contrast (Fischer et al., 2013). This suggests visual processing at the saccade target likely starts prior to fixation onset and makes the current findings hard to reconcile with purely post-saccade processing.

Effects of pre-saccadic stimulus on post-saccadic scalp potentials have been demonstrated in a variety of studies. For instance, the presence or spatial frequency of simple stimuli before a saccade is reflected in an occipital positivity 50-75 ms after fixation onset (Bellebaum & Daum, 2006; Kazai & Yagi, 1999). These have been interpreted as signals related to the updating of retinal location (Peterburs et al., 2011) leveraging an efference copy, consistent with a saccade-lock of this early contrast.

More recent work emphasizes predictive processing based on the pre-saccadic stimulus. For instance, if a pre-saccade stimulus changes during a saccade, occipital electrodes show differences starting at 200 ms after the saccade (Ehinger et al., 2015). Similarly, during reading, upcoming words that appear in the periphery affect occipital electrodes when they are fixated after a saccade (Dimigen et al., 2012). A recent study on face perception shows that a scrambled face presented in the periphery prior to fixation decreases responses starting at 180ms post-fixation (Buonocore et al., 2020). The picture that emerges from these studies is that the pre-saccadic stimulus primes visual processing after the subsequent saccade. These experiments were all conducted with highly constrained stimuli and tasks. Here, with an unconstrained detection task and relatively naturalistic dynamic stimulus, we find both an early effect of the pre-saccade target location, which may relate to updating, as well as a later response likely associated with surprise, which may relate to predictive processing.

One caveat to our conclusions is that the lambda complex is modulated by saccade amplitude (Yagi, 1979). The method we used to detect saccades based on EOG had the advantage that it detected saccades of approximately the same duration, and hence, size (Bahill et al., 1975).

We generally found no differences in saccade duration in all the contrasts we performed. We did however find differences in saccade-locked TRFs starting with saccades onset, in the contrast between peripheral and central targets. This is likely the result of difference in saccade dynamics. However, given the lack of a difference in saccade duration, it is unlikely that amplitude differences after the saccade (at 129ms) are the result of altered saccade dynamics. One caveat to our study is that we were not able to detect and analyze smaller saccades, which may have differed between these two conditions.

While much of the literature on the lambda complex is locked to fixation onset, here we favored the saccade onset because it is more clearly defined in time in the EOG, which we use to detect saccades and fixations. In general, saccade-related signals are more time accurate, as compared to fixation-related signals, e.g. (Katz et al., 2020). Nevertheless, given the close correspondence of the saccade and fixation locked lambda (Thickbroom et al., 1991), we expect however that most results would remain if we had performed a fixation-locked analysis. We recommend recording eye position with sufficient temporal and spatial resolution to detect saccades onset and offset, to overcome the limitation of the present study.

## Conclusion

The literature suggests a continuity of visual perception during saccades, with processing starting prior, an inhibition during, and an enhancement after the saccade. This existing work largely used discrete stimuli and constrained eye movements. Here we used a target detection task embedded in a video game to provide a more naturalistic setting with free-viewing. We find that the presence of the target, its visibility and its location affect neural responses early after the saccade. Overall we conclude that during natural viewing of dynamic scenes, neural processing of the visual stimuli, including orienting, start before a saccade and continue throughout the saccade, apparently unencumbered by saccadic inhibition. To our knowledge, this work is the first to report continuity of visual processing throughout the saccade during visual search in a realistic, free-viewing environment.

Methodologically, the results of the present study steer the conversation away from visual evoked responses locked to the timing of stimulus presentation and towards a focus on eye movements and their essential relation to task engagement, which motivates where we look.

## Methods

### Data collection and procedure

Healthy, right-handed male subjects between the ages of 20 and 40 (mean 28.3) were recruited among ARL and other government employees. Subjects gave informed consent in accordance with the requirements of the Institutional Review Board of the ARL. The experiment was conducted in an electrically-shielded, sound-dampened room. Prior to the primary experimental task, subjects performed an eye-calibration (see below). For the primary experimental task, subjects were driven around a simulated 3D environment and were required to discriminate human entities as either a non-target (an image of a man) or a target (an image of a man with a gun) and tables as either a non-target (an image of a table with a clear view under it) or a target (an image of a table with an obstructed view of the space under it). Stimuli were presented for 1 second with an inter-stimulus interval (ISI) of 2 ± 0.5 s. For each target the subject responded by pushing a button with the left or right hand to indicate the target type. The button used to indicate target type (left or right) was counterbalanced across subjects in an alternating fashion. Participants were instructed to use the right index finger for pressing the right button and the left index finger for the left button.

### Experimental stimuli and task description

The subject was presented with 300 human and 300 table targets. These 600 targets were split into two successive runs. In other words, during the first 15-minute block of the experiment, the subjects encountered targets 1-300. During the second 15-minute block the subjects encountered targets 301-600. The location and placement of targets were randomly predefined and were held consistent across all subjects. In other words, each subject saw the same target at the same location. The order of presentation of 15-minute blocks was counterbalanced across subjects. The location in which targets appeared was visually verified during pilot testing to be at logical locations, i.e. level with the ground plane and with an unobstructed line of sight.

During each block visibility conditions were altered through the use of a fog-like overlay on the image. Throughout each block the fog would appear for a few seconds, up to a minute, and then disappear.

Finally, during each 15 minute block, subject scores were presented on the bottom of the screen. Subjects were instructed to try to beat a virtual competitor. Unbeknownst to the subject, the virtual competitor’s score was adjusted to follow the subject’s score with extended time frames when the participant was winning or losing. In the easy condition, this difference was such that the player was winning most of the time including at the end. During the hard condition, the virtual competitor was winning about half the time and would often win at the end.

### Equipment

The EEG configuration was a 64-Channel Biosemi Active II (Amsterdam, The Netherlands) recorded at 1024Hz. Online referencing was done to the Common Mode Sense electrode which was later offline re-referenced to the mastoid electrodes. Eye tracking was done using a 60Hz eye-tracker faceLAB 4.2 by Seeing Machines (Fystick, Australia). External electrodes from the BioSemi system were placed on both mastoids for signal reference, next to the outer canthus of each eye to record horizontal EOG, and directly below the left eye in the inferior orbital margin to obtain a differential vertical EOG record when paired with the signal from the Fp1 electrode in the standard 64 channel montage. Additional external electrodes were placed at the top and bottom of the left masseter jaw muscle, and below the left breast to obtain a basic ECG waveform.

### EEG Preprocessing

EEG was filtered with a 0.3 to 250Hz bandpass filter and eye-movement artifacts corrected by regressing out a single left vertical and a single right horizontal electrooculogram (EOG) channels and two left frontal channels (Fp1 and AF7; for display purposes Fp1 and AF7 were replaced with their right side mirror opposites Fp2 and AF8). Subsequently we used robust principal component analysis (Candès et al., 2011), which implicitly removes outlier channels and myographic activity. EEG was downsampled to 256Hz for TRF analysis. All data preprocessing and analysis were implemented in MATLAB.

### Gaze position

Eye tracking data was used to determine target location relative to gaze position. For this purpose raw eye tracking data (60Hz sampling frequency) were calibrated to the 1024 by 768 pixel space of the screen by using the calibration tasks prior to the main experiment. Then they are asked to look at dots that appear in random sequence at the four corners or the center of the screen and click with the mouse when the dot appears. The gaze coordinates at the time of the mouse clicks were used to create an affine transform that maps the gaze position to screen coordinates of the corresponding dots. To reduce variance in the gaze position we applied a temporal filter (rectangular window of size 0.25s). The result was validated using gaze position data collected during a smooth-pursuit task (follow a slowly moving ball on the screen). The screen size was 24’’ (51.1 × 29.9 cm) and subjects were seated approximately 60 cm from the screen resulting in a visual stimulus that covers approximately 44.2° by 33.9° in visual angle.

### Saccade detection

Saccades onset times and duration were detected from the electrooculogram because recordings of the eye position did not have sufficient temporal resolution after temporal smoothing. Specifically, we used a saccade detection algorithm based on EOG activity (Toivanen et al., 2015) and applied it here to the raw EOG signal (horizontal minus vertical channels) at a resolution of 1024Hz.

### Data Analysis

Temporal response functions (TRFs) were used to derive overlapping evoked responses using a conventional linear system identification approach. This approach assumes that all the responses during the target trials are a linear superposition of 5 types of events. The presentation of the target, button responses, and saccades, which were further divided into those following the target (100 and 450ms), those within 1s of target presentation but outside this range, and spontaneous saccades, which are more than 1s away from the target. Each type of response has a stereotypic temporal response filter (TRF, equivalent to an impulse response) that when convolved with all the onsets of each respective event and added together across event types will produce an estimate of EEG signal. Algebraically the convolution is constructed by the matrix multiplication:

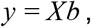

where *X* is a binary Toeplitz matrix indicating when events happen, *b* are the TRFs, and *y* is the estimated EEG time series. The Toeplitz matrix implements the convolution and can be concatenated to form a block-Toepliz matrix that implements the sum over the five event types:

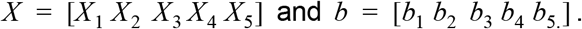

To derive the TRFs we use conventional least-squares optimization:

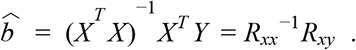

*R*_*xx*_ and *R*_*xy*_ are the auto- and cross-correlation matrices of *x* and *y*. *R*_*xy*_ captures the conventional trial-averaged evoked responses. The inverse of *R_xx_* removes correlation induced but correlated events and responses that overlap in time. For memory efficiency the construction and multiplication of *X* are done using sparse matrix form in MATLAB before switching back to full matrix form for *R*_*xx*_ and *R*_*xy*_. For further memory efficiency the TRFs were computed for each subject individually and then averaged across subjects to give the group estimate TRF. Before averaging across subjects, each TRF was baseline corrected using the 125ms before onset.

The approach used here is essentially a classic multiple-input single-output (MISO) systems identification approach. To our knowledge it has been first used for EEG in the context of continuous stimuli driving evoked responses (Crosse et al., 2016; Lalor et al., 2006). In contrast to this previous work we found no need to regularize the inverse of *R_xx_*, which is sometimes done to reduce noise. The same approach has been also used to disentangle overlapping responses of consecutive eye movements (Dandekar et al., 2011; Dimigen & Ehinger, 2021; Guérin-Dugué et al., 2018). The work of Dimigen & Ehinger extend the approach by including classic interaction terms to determine if different types of events influence each other (multiplicatively). In that work a TRF is estimated for each type of event and a separate TRF is estimated to capture the difference between conditions. In contrast, here we estimate a TRF for each condition separately and then take a difference. Given the linearity of the model, these approaches are equivalent (except for the interaction terms, which we do not use here).

For the analysis of spontaneous vs target-evoked saccades we estimated a single model *b* using the data from the clear condition and fog condition. For the comparison of fog and clear conditions a separate model *b* was estimated for each condition. For the comparison of central and peripheral targets we estimate a separate model *b* for each type of saccades.

Statistical significance of contrast in the TRFs between different conditions were determined using a permutation test. 500 surrogate contrasts were created by shuffling trials between conditions. This was done for the contrasts in saccade type (Fig. 3), target visibility (Fig. 4), target location (Fig. 5), and target threat type (Fig. S7). To compare across game difficulty 500 surrogates were created by shuffling subjects between the two game difficulties (not showns). Across 64 channels and 320 time points (spanning −250 to 1000 of the trial) there were 20480 individual tests that were FDR corrected (Benjamini & Hochberg, 1995) with an alpha of 0.05.

## Acknowledgments

Funding source: ARL/DSO W911NF-10-2-0022 (Subaward from DCS Corp: APX02-N013).

