## Supplemental Figures for "During natural viewing, neural processing of visual targets continues throughout saccades"

**Supplement**

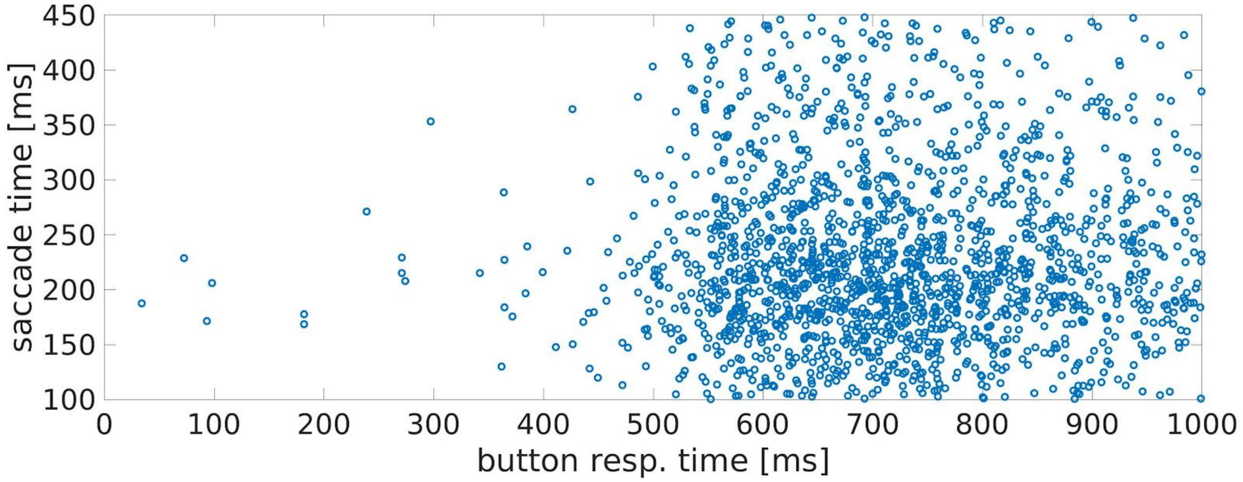


**Fig. S1: Button press supplement: A:** Button press TRF **B:** Scatter plot of all saccades to target times plotted against button response times. There showing no relationship between when subjects looked towards a target and how fast they responded to it.

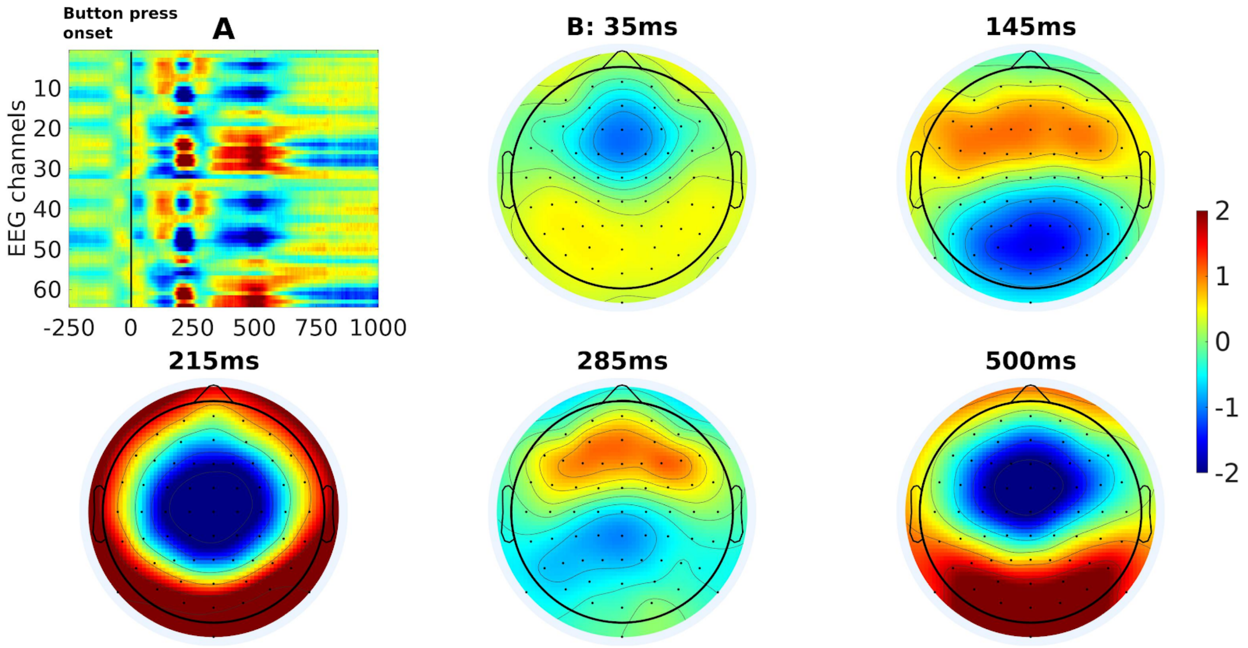


**Fig S2: Button press response TRF A:** Response across all 64 channels locked to button press at 0ms. **B:** Representative topographic plots of post-button press peaks at 35, 145, 215, 285 and 500ms.

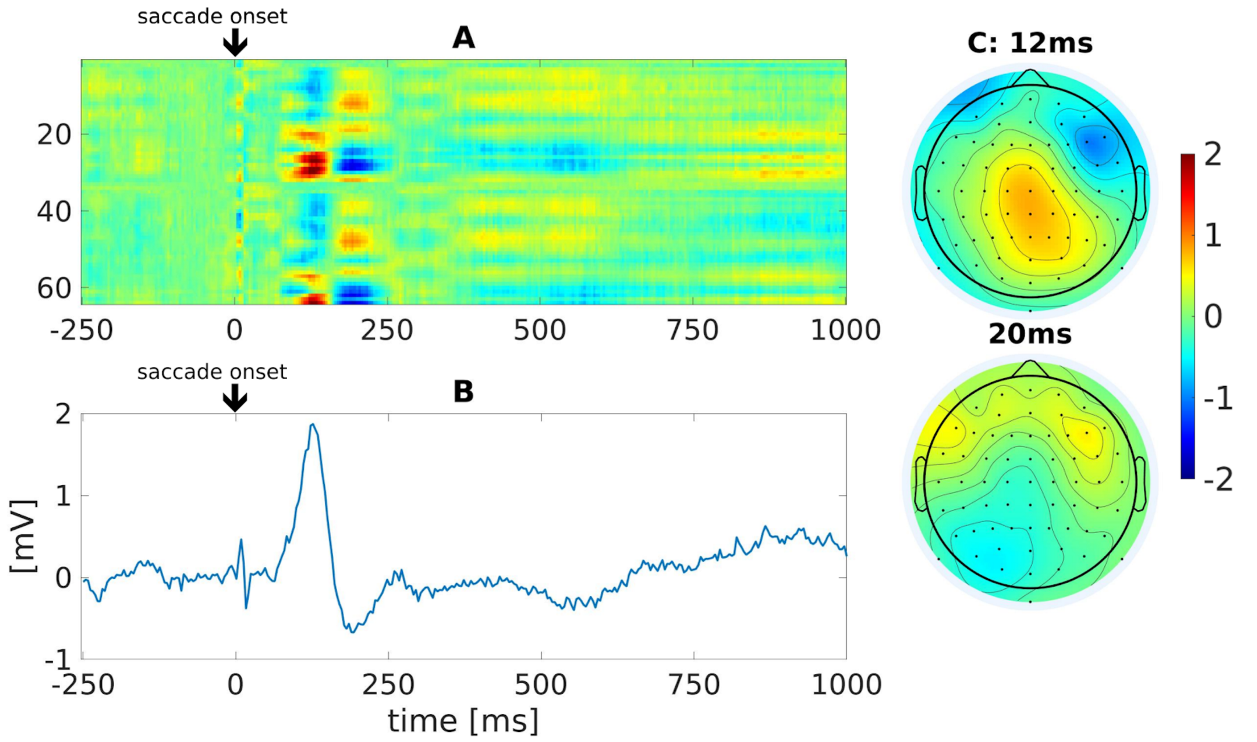


**Fig. S3: Saccade-locked TRF of all targets showing the stereotypical lambda complex A:** TRF locked to target presentations (demarcated by 0 ms). **B:** Average trace of electrodes CPz, POz, and Oz. **C:** Respective topographic snapshot of peaks at 12ms and 20ms. Peak at 12ms is indicative of the spike potential of the saccadic lambda complex.

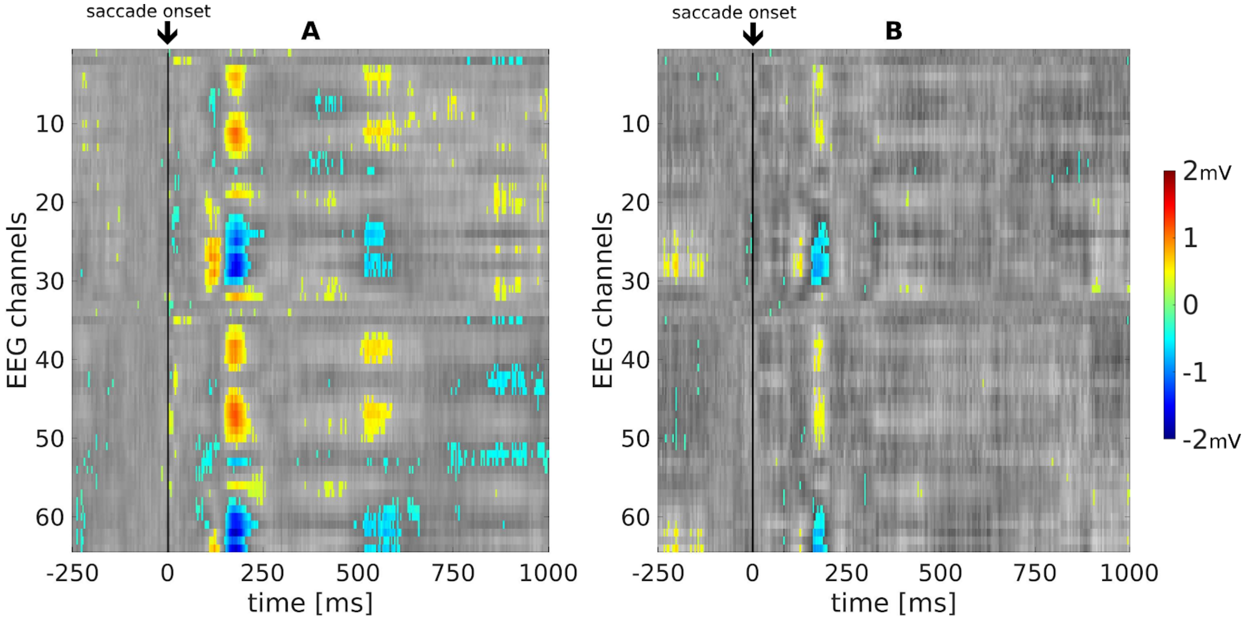

**Fig. S4: Target-elicited saccades and spontaneous saccades contrast A:** Saccade-locked TRF shows a replicable difference for both "easy" and (target: N=1749, spontaneious: N=12437) **B:** "hard" game conditions (target: N=1774, spontaneous: N=13937) .

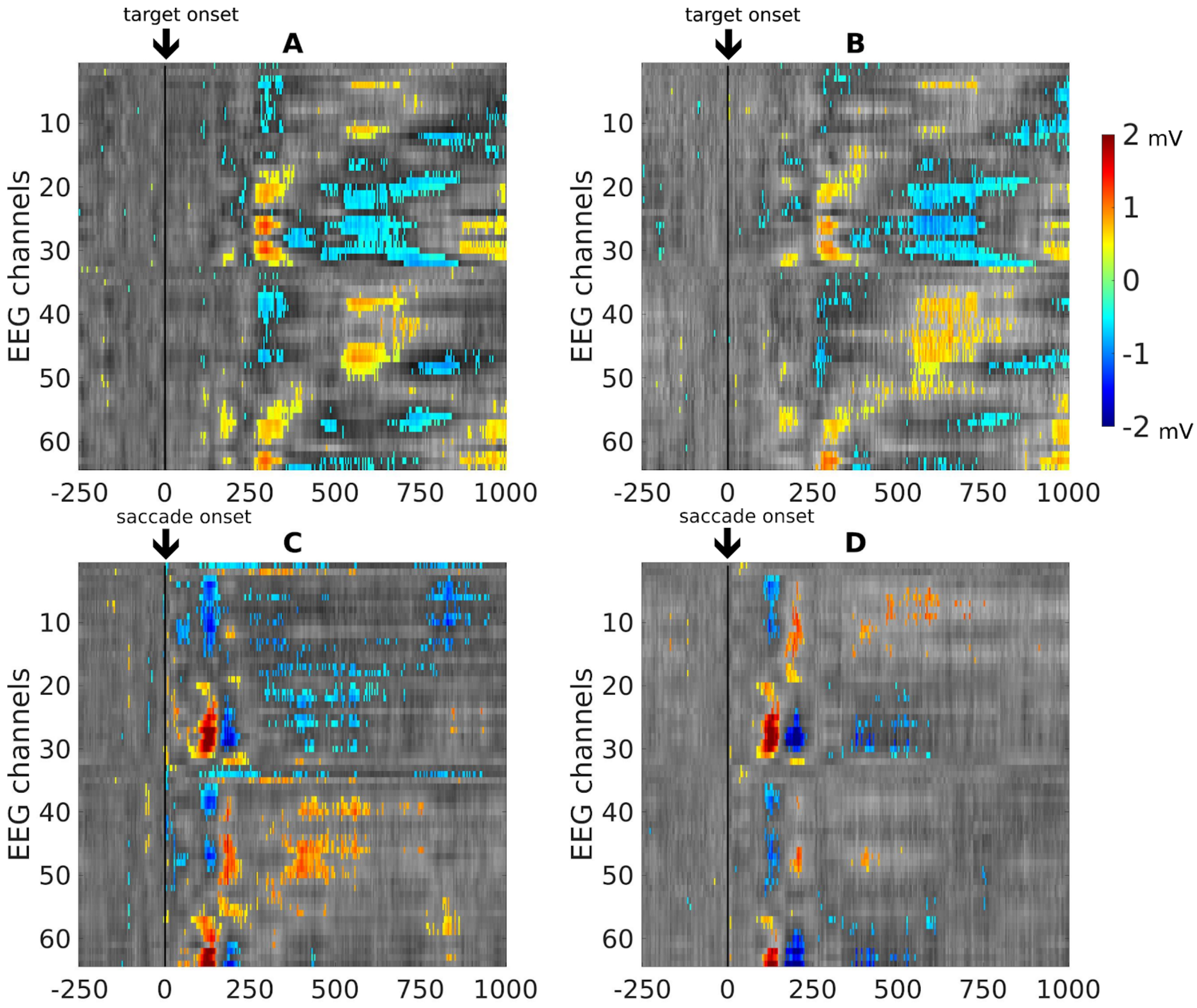


**Fig. S5: Target visibility supplement: A:** Target-locked TRF differences between clear and  for the "easy" game condition replicates with the (clear: N=3159, fog: N=876) **B:** target-locked "hard" game condition (clear: N=2784, fog: N=619). **C:** Similarly the saccade-locked TRF shows a replicable difference for both "easy" and (clear: N=1382, fog: N=367) **D:** "hard" game conditions (clear: N=1461, fog: N=313) .

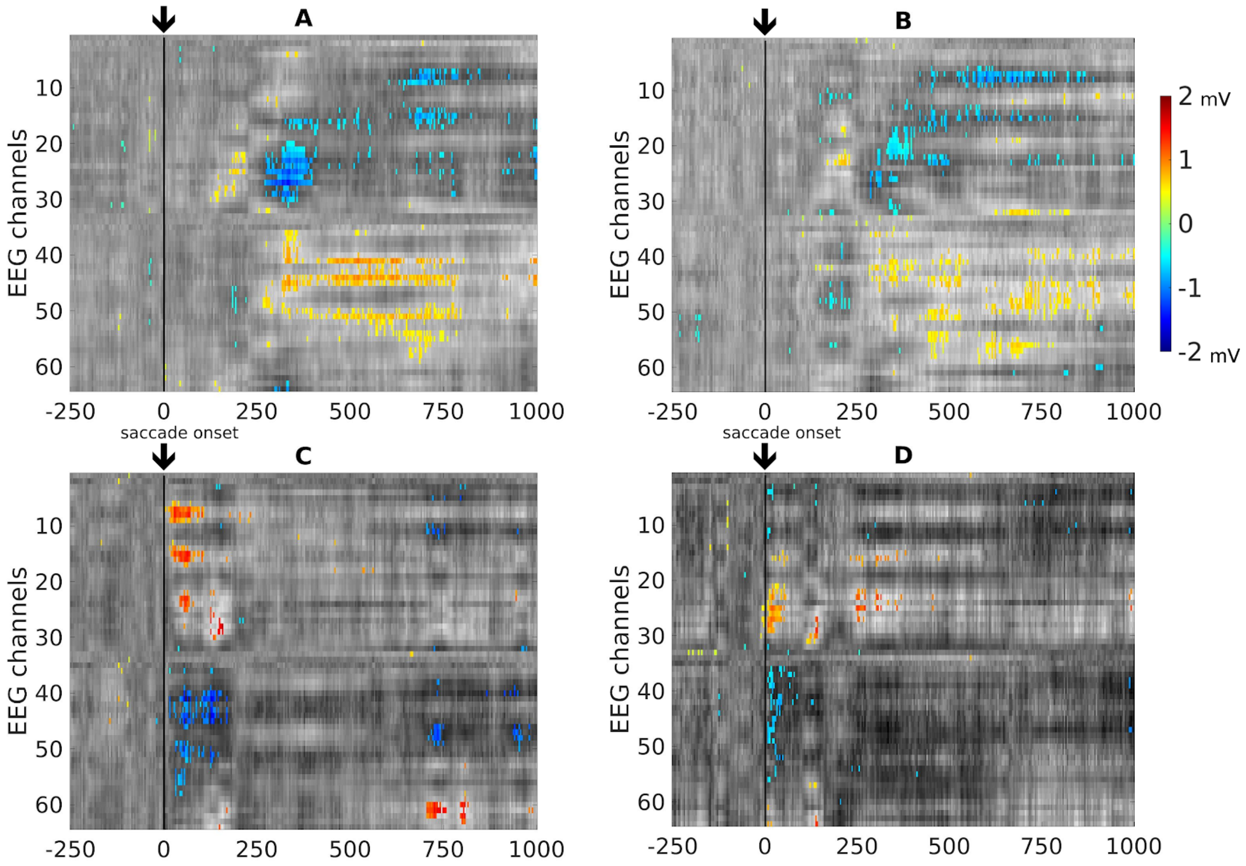


**Fig. S6: Target location supplement: A:** Target-locked TRF differences for shows a strong late contrast between peripheral and central for the “easy” game condition (peripheral: N=1546, central: N=1613) which replicates with the **B:** target-locked “hard” game condition (peripheral: N=1310, central: N=1474) . **C:** Saccade-locked TRF shows a replicable difference for both “easy” (peripheral: N=830, central: N=552) and **D:** “hard” (peripheral: N=838, central: N=623) game condition.

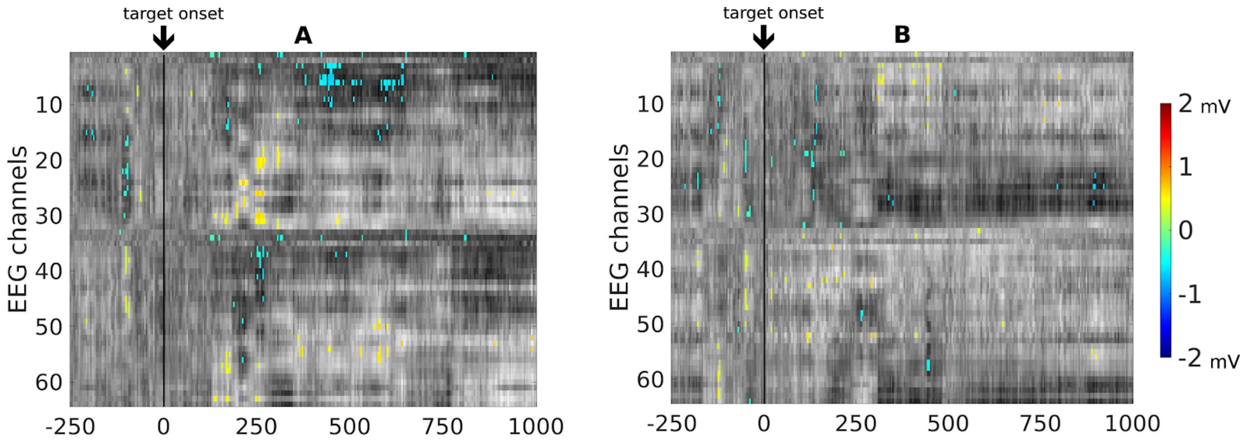

**Fig S7: Threat contrast supplement: A:** “easy” game condition (threat: N=1571; no-threat: N=1588) showing significance between 175-250ms and 430-645 but this does not replicate for the **B:** “hard” game conditions (threat:  N=1328; no-threat: N=1456). Saccade-locked results showed no significance.
